# A probabilistic model for indel evolution: differentiating insertions from deletions

**DOI:** 10.1101/2020.11.22.393108

**Authors:** Gil Loewenthal, Dana Rapoport, Oren Avram, Asher Moshe, Alon Itzkovitch, Omer Israeli, Dana Azouri, Reed A. Cartwright, Itay Mayrose, Tal Pupko

**Affiliations:** The Shmunis School of Biomedicine and Cancer Research, George S. Wise Faculty of Life Sciences, Tel Aviv University, Tel Aviv 69978, Israel; School of Plant Sciences and Food Security, George S. Wise Faculty of Life Sciences, Tel Aviv University, Tel Aviv 69978, Israel; The Biodesign Institute, Arizona State University, Tempe, Arizona, USA; School of Life Sciences, Arizona State University, Tempe, Arizona, USA

**Keywords:** molecular evolution, evolutionary models, indels, alignments, approximate Bayesian computation, Neural networks

## Abstract

Insertions and deletions (indels) are common molecular evolutionary events. However, probabilistic models for indel evolution are under-developed due to their computational complexity. Here we introduce several improvements to indel modeling: (1) while previous models for indel evolution assumed that the rates and length distributions of insertions and deletions are equal, here, we propose a richer model that explicitly distinguishes between the two; (2) We introduce numerous summary statistics that allow Approximate Bayesian Computation (ABC) based parameter estimation; (3) We develop a neural-network model-selection scheme to test whether the richer model better fits biological data compared to the simpler model. Our analyses suggest that both our inference scheme and the model-selection procedure achieve high accuracy on simulated data. We further demonstrate that our proposed indel model better fits a large number of empirical datasets and that, for the majority of these datasets, the deletion rate is higher than the insertion rate. Finally, we demonstrate that indel rates are negatively correlated to the effective population size across various phylogenomic clades.

## Introduction

Insertions and deletions (indels) shape genes and genomes and are fundamental in molecular evolution research (Cartwright, 2009). Indels are of great importance for ancestral sequence reconstruction (Ashkenazy et al., 2012; Vialle, Tamuri, & Goldman, 2018), and substantially contribute to divergence among species (Anzai et al., 2003; Britten, 2002; Britten et al., 2003; Wetterbom et al., 2006). Fitch (1973) was the first to observe that deletions may be more common than insertions, however, this observation was based on very few protein sequences. De Jong and Ryden (1981) analyzed a much larger set of proteins, and suggested that deletions are four fold more frequent than insertions, and that this phenomenon is an inherent property of the replication mechanism. In support for this hypothesis, Graur et al. (1989) found over three times more deletions than insertions in processed human and rodent pseudogenes, suggesting that it is mutations rather than selection that drive the excess of deletion over insertion events. This deletion bias was confirmed by numerous other studies (Kuo & Ochman, 2009; Mira et al., 2001; Ogata et al., 1996; Ophir & Graur, 1997; Petrov et al., 1996; Fan et al., 2007; Van Passel et al., 2007; Zhang & Gerstein, 2003).

Regarding the distribution of indel length, it was repeatedly observed that in both proteins and DNA sequences, single-site indels are the most frequent and the occurrences of indels decline monotonically as a function of their length (Benner, Cohen, & Gonnet, 1993; Golenberg, Clegg, Durbin, Doebley, & Ma, 1993; Gu & Li, 1995; Pascarella & Argos, 1992; Qian & Goldstein, 2001). Two distributions were proposed for the indel length: geometric and Zipfian. It was previously shown that the Zipfian distribution better fit biological datasets, both for proteins (Benner et al., 1993) and for non-coding regions (Saitou & Ueda, 1994). Gu and Li (1995) found only small differences in the size distribution of deletions and insertions. When insertions and deletions were treated together, the parameter of the Zipfian length distribution varied from 1.70 in primate globin non-coding regions to 1.93 in non-coding mitochondrial DNA. Of note, these early studies were based on small datasets, such that only a few indel events were considered. In another study that analyzed coding and non-coding indels among 18 mammalian genomes, ,differences were found both among species and between insertions and deletions: the Zipfian parameter ranged from 1.059 to 1.883, when modeling the length distribution of deletions in chimpanzee to insertions in rabbit, respectively (Fan et al., 2007). In all these studies, the indel parameters were inferred based on indel counts, and thus only indels which could be reliably aligned among the analyzed sequences were included.

Probabilistic-based models for indels are far less developed compared to substitution models. This might be the case since indel models violate the assumption of site independence, which complicates the computation of the likelihood function (Cartwright, 2005; Fletcher & Yang, 2009). More elaborate methodologies to estimate indel parameters include Cartwright’s lambda.pl Perl script released with the DAWG simulation package (Cartwright, 2005). It assumes a Poisson distribution for indel rates and estimates the distribution using the maximum likelihood paradigm. The method uses linear regression to find the best fitted Zipfian distribution for the indel length and takes the average length of the input sequences as the root length. Two additional methods are based on Hidden Markov Model (HMM) between pairs of divergent sequences (Cartwright, 2009; Lunter, 2007). In Lunter (2007), biases introduced by alignment programs, such as ‘gap attraction’, the tendency of alignment algorithms to merge two independent gaps, were explicitly accounted for and gap lengths were assumed to follow a mixture of geometric distributions. Cartwright (2009) used expectation maximization algorithm based on a pairwise HMM for the inference of model parameters. This method assumes independence between indel events and ignores overlapping indels. These methods were restricted to pairwise sequences, and thus could not distinguish between insertion and deletion rates.

We have previously developed SPARTA (Karin et al., 2015), a simulation-based algorithm to learn indel parameters from input MSAs. SPARTA is an ad-hoc methodology that is not rooted in probabilistic theory. We later developed SPARTA-ABC (Karin et al., 2017), which is based on the Approximate Bayesian Computing (ABC) methodology, a statistically rigorous methodology for the inference of model parameters. The ABC framework, first introduced in molecular evolutionary studies for population genetics (Beaumont et al., 2002), has been utilized successfully to estimate parameters in complex models, in which the likelihood function is challenging to compute. ABC was successfully employed, for example, for estimation of the effective population size from a sample of microsatellite genotypes (Tallmon et al., 2008), estimation of divergence times and admixture by analyzing whole genomes of chimpanzee and bonobo populations (Kuhlwilm et al., 2019), and inference of relevant parameters relating to selective sweeps; that is, selection coefficient, time of selection onset, recombination rate and mutation rate at neutral loci (Przeworski, 2003).

The underlying indel probabilistic model in SpartaABC assumes that the insertion rate (number of insertions events per substitution event) equals the deletion rate. It further assumes that the length of an insertion event (number of newly introduced nucleotides or amino acids) has the exact same distribution as the length of a deletion event. As stated above, these assumptions are known to be an oversimplification of indel dynamics. In this study, we develop a more realistic alternative by assigning different parameters for insertions and deletions. We also compare two model-selection schemes to determine whether the richer model better describes indel evolutionary dynamics compared to the simple model: a classic ABC approach in which the estimated posterior distribution of each model is approximated by the relative frequency of simulations generated from each model (Pritchard et al., 1999), as opposed to a newly developed neural-network-based-approach, which was previously developed in the field of population genetics (Mondal et al., 2019). We show that the latter model selection scheme is more accurate than the previous one. Comparing these two model-selection schemes enabled us to demonstrate that the richer model fits a large number of empirical biological datasets, lending further statistical support for the hypothesis that deletion dynamics are more rampant than those of insertions. Finally, we studied how indel evolutionary dynamics varies among different phylogenetic groups. This analysis allowed us to confirm previous results from Sung et al. (2016), who demonstrated a negative correlation between indel rate and effective population size. This negative correlation corroborates the drift barrier hypothesis, i.e., natural selection is limited by the extent of drift, whose intensity is inversely proportional to the effective population size (Sung et al., 2016; Sung et al., 2012).

## Results

### Indel models

We describe two indel models, a simple indel model (SIM) and a rich indel model (RIM), which alleviates some of the assumptions made in SIM. The parameters of both models are summarized in table 1. In SIM, insertions and deletions are assumed to have the same rates and length distributions. Thus, SIM has three parameters: (1) Indel-to-substitution-rate ratio (*R*_*ID*), note that this parameter quantifies the sum of the insertion and the deletion rates, which are assumed to be equal in this model; (2) The insertion length distribution parameter (*A*_*ID*), which dictates the distribution of the lengths of newly inserted or deleted segments. Qian & Goldstein (2001) showed that the frequencies of indels that are several dozens of amino acid long are lower than their expected frequencies, when the expectation is computed based on the length distribution of shorter indels. Thus, in our models, it is assumed that this length is distributed as truncated Zipfian (power-law) with maximum indel size of 50 amino acids and a rate parameter *A*_*ID* (*A*_*ID* stands for the “*a*” parameter of the Zipfian distribution for insertion/deletion length). (3) The sequence length at the root of the tree (*RL*). In RIM, different indel parameters are assigned to insertions and deletions, resulting in five free parameters. In addition to the root length, two parameters dictate the indel rates, one for insertions (*R*_*I*) and one for deletions (*R*_*D*). Similarly, two “*a*” parameters are assumed, one dictating size distribution for insertions (*A*_*I*) and one for deletions (*A*_*D*).

**Table 1.** The Simple Indel Model (SIM) and Rich Indel Model (RIM) parameters and their description.

| Model | Parameter name | Description |
| --- | --- | --- |
| *SIM, RIM* | RL | The sequence length at the root of the tree |
| *SIM* | R_ID | Indel-to-substitution-rate ratio |
|  | A_ID | Indel length distribution parameter |
| *RIM* | R_I | Insertion-to-substitution-rate ratio |
|  | R_D | Deletion-to-substitution-rate ratio |
|  | A_I | Insertion length distribution parameter |
|  | A_D | Deletion length distribution parameter |

### Prior distributions of model parameters

Model parameters are inferred using ABC. In this Bayesian inference scheme, prior distributions over model parameters have to be chosen. We assume the following prior distribution: (1) The indel to substitution rates are assumed to be uniformly distributed in the range [0, 0.1] for R_ID (SIM) and [0, 0.05] for R_I and R_D (RIM). (2) The parameters that dictate the indel length distribution (*A*_*ID* for SIM, and *A*_*I* and *A*_*D* for RIM), are assumed to be uniformly distributed in the range [1.001, 2]. (3) The *RL* parameter range is determined according to the input sequences, as follows: let *l*_*s*_, *l*_*l*_, be the length or the shortest and longest sequences, respectively, of the unaligned sequences, then the range of *RL* is assumed to be uniformly distributed in the range [0.8*l*_*s*_,1.1*l*_*l*_]. We note that increasing the range of the prior distributions had little effect on the results (not shown).

### Inference outline

The ABC inference scheme relies on several components/steps: (1) Generating simulations; (2) Computing summary statistics; (3) Assigning summary statistics weights; (4) Accepting a subset of the simulations; and (5) Inferring the posterior distributions and point estimates. These components are described in detail below. Here, we first present a general outline of the algorithm. The input required to infer the model parameters for a dataset in question is a set of multiple sequence alignment (MSA) and its associated (rooted) phylogenetic tree, including the topology and its associated branch lengths. Next, a large set of simulated MSAs is generated based on model parameters sampled from the prior along the input phylogenetic tree. Next, summary statistics are computed for both the input MSA and each of the simulated MSAs. Summary statistics weights are next computed from a subset of these simulations and are then used to compute distances between the summary statistics of the input MSA and each of the simulated MSAs. A small subset of simulations, for which the distance is very small, is kept. Intuitively, the kept simulations resemble the input data in terms of indel dynamics and can be used to get a point estimate of the model parameters of the dataset in question. The distribution of model parameters used to generate this subset is a good approximation for their posterior distribution (Sisson, 2018). Thus, the last step of the algorithm is to infer posterior distribution and point estimate for all model parameters.

### Simulator

Existing tools for simulating sequences such as DAWG 2.0 (Cartwright, 2005) and INDELible (Fletcher & Yang, 2009) account for both substitution and indel events. For the purpose of inferring the relevant summary statistics, the information regarding substitutions can be ignored. Thus, simulations can be performed without substitutions, thereby reducing simulation running times, which are a major component of the ABC inference scheme. To this end, we developed an indel simulator for SIM and RIM that ignores substitution events.

### Summary statistics

The 27 summary statistics calculated in the inference scheme are described in table 2. This list extends the 11 summary statistics previously used by Karin et al. (2017) and also includes summary statistics that can help to differentiate insertion from deletion events. For example, the 13^th^ summary statistic, i.e., number of MSA columns that contain a single gap, provides information on deletion rates, as a column with a single gap typically reflects a single deletion event. Another example is the 17^th^ summary statistic, which counts the number of MSA columns in which a single residue gap is found in all but one sequence. Such a column, most likely, reflects an insertion of a single residue in a branch leading to a leaf of the tree. Notably, such a column may result from a deletion event as well. The ABC approach does not assume that this is certainly an insertion event, but rather, all summary statistics are considered together and their values provide information regarding the posterior probability of the model parameters.

**Table 2.** The 27 summary statistics used in the ABC scheme.

| # | Summary statistic |
| --- | --- |
| 1 | Total number of gap blocks in the alignment |
| 2 | Total number of unique gap blocks in the alignment |
| 3 | Average gap block length |
| 4 | Average unique gap block length |
| 5 | Number gap blocks of length one |
| 6 | Number gap blocks of length two |
| 7 | Number gap blocks of length three |
| 8 | Number gap blocks of length four or more |
| 9 | Alignment length |
| 10 | Minimum length of sequence in the alignment |
| 11 | Maximum length of sequence in the alignment |
| 12 | Number of MSA columns with zero gaps |
| 13 | Number of MSA columns with one gap |
| 14 | Number of MSA columns with two gaps |
| 15 | Number of MSA columns with $n - 1$ gaps |
| 16 | Number of gaps of length one in one sequence |
| 17 | Number of gaps of length one in two sequences |
| 18 | Number of gaps of length one in $n - 1$ sequences |
| 19 | Number of gaps of length two in one sequence |
| 20 | Number of gaps of length two in two sequences |
| 21 | Number of gaps of length two in $n - 1$ sequences |
| 22 | Number of gaps of length three in one sequence |
| 23 | Number of gaps of length three in two sequences |
| 24 | Number of gaps of length three in $n - 1$ sequences |
| 25 | Number of gaps of length at least four in one sequence |
| 26 | Number of gaps of length at least four in two sequences |
| 27 | Number of gaps of length at least four in $n - 1$ sequences |

### Computing weights for the summary statistics

Let *D*_*i*_ and *D*_*s*_ denote an input MSA and a simulated MSA, respectively. Let *S*(*D*_*i*_) and *S*(*D*_*s*_) be summary statistic vectors associated with *D*_*i*_ and *D*_*s*_, respectively. In order to decide whether or not to keep a simulation, a weighted Euclidean distance is computed between *S*(*D*_*i*_) and *S*(*D*_*s*_) as follows:

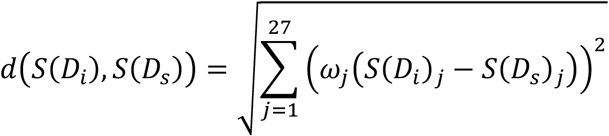

where the subscript *j* is the summary statistic index and *ω*_*j*_ denotes the weight of the *j*^*th*^ summary statistic. The various summary statistics differ in their magnitude, so different weights are required to ensure that all the summary statistics contribute approximately equally to the distance. Hence, the weight of each summary statistic is set as 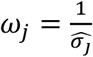, where 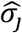 is the estimated standard deviation of the *j*^*th*^ summary statistic across *B* burn-in simulations with indel parameter values drawn at random from the prior. We set *B* = 10,000 because for this value, the vector of weights practically converged in all cases (not shown).

### Acceptance/rejection criterion

The weighted Euclidian distance is calculated for N_s_ simulations. By default, N_s_ = 100,000 (using 1,000,000 simulations did not significantly improve the performance; not shown). The set of accepted simulations are chosen such that the rate of accepted simulations is *p* of the total simulations (Beaumont et al., 2002). In this study the *p* parameter was set to 0.1% (100/100,000) of the simulations (0.1% yielded the best performance in a small-scale simulation study, not shown).

### Inference of posterior distributions and point estimates

The posterior distribution is approximated by the distribution of the parameter values of the accepted simulations. In the classic ABC framework, point estimates for model parameters are obtained by averaging the parameters of the accepted simulations (Tavaré et al., 1997). It was previously shown that alternative methodologies, such as regression, can yield more accurate point estimates (Beaumont et al., 2002; Bertorelle et al., 2010). Blum et al. (2013) have previously suggested to combine machine-learning algorithms and ABC for the accurate inference of model parameters. In this work, we test the applicability of machine-learning regression models for inferring indel parameters. To this end, we compared three approaches: a classic ABC approach in which the parameters of the accepted simulations are averaged (Beaumont et al., 2002); and two machine-learning algorithms named Ridge regression (Hoerl & Kennard, 1970) and Lasso regression (Tibshirani, 1996). In the machine-learning approaches, we train a regression model for each model parameter. The training data are the accepted simulations and each training data point is a vector of summary statistics (normalized to have zero mean and standard deviation of one) and its associated known parameter, i.e., the parameter that was used to simulate the training data. The regularization parameter for the machine-learning algorithms is tuned per indel parameter by a 10-fold cross validation procedure. To infer each model parameter of an empirical data, the vector of summary statistics is given as input to the corresponding trained regression model.

### Inference accuracy on simulated data

We tested the accuracy of SpartaABC in inferring model parameters by simulating datasets with model parameters sampled from the prior, based on a specific tree topology and an MSA sampled from the EggNOG database (Huerta-Cepas et al., 2019). The MSA contains seven sequences, with a mean sequence length of 1,384 amino acids. As described in the section ‘inference of posterior distributions and point estimates’, we tested three different inference procedures. To quantify inference accuracy, we computed the *R*^2^ values between the true parameters and the inferred ones, over 200 random different parameter combinations sampled from the prior distribution. The obtained *R*^2^ values for the Lasso regression were 0.92, 0.97, 0.96, 0.81, and 0.83 for *RL*, *R*_*I*, *R*_*D*, *A*_*I*, and *A*_*D*, respectively (fig. 1). Similar results were obtained with Ridge regression and lower *R*^2^ values were obtained for the averaging-based inference (supplementary table S1, Supplementary Material online). We extended this simulation analysis, repeating the simulation scheme for 12 additional datasets that differ from the one presented in figure 1 with respect to tree topologies, total branch lengths, number of species, and sequence length (supplementary table S2, Supplementary Material online). These simulations demonstrate that the estimates of the parameters controlling the indel rates and root length (*R*_*I*, *R*_*D* and *RL*) are more accurate than those dictating the length distribution of indels (*A*_*I* and *A*_*D*). As expected, the inference accuracy strongly depends on total branch lengths (supplementary fig. S1, Supplementary Material online). Our results further suggest that SpartaABC provides relatively unbiased estimates for all model parameters, as the slope of the regression fit was not significantly different from 1.0 (supplementary table S2, Supplementary Material online). We conclude that SpartaABC provides accurate estimates of model parameters, most notably for the indel rates and the root length, as long as sufficient indels have accumulated to allow reliable inference.

**FIG. 1.**
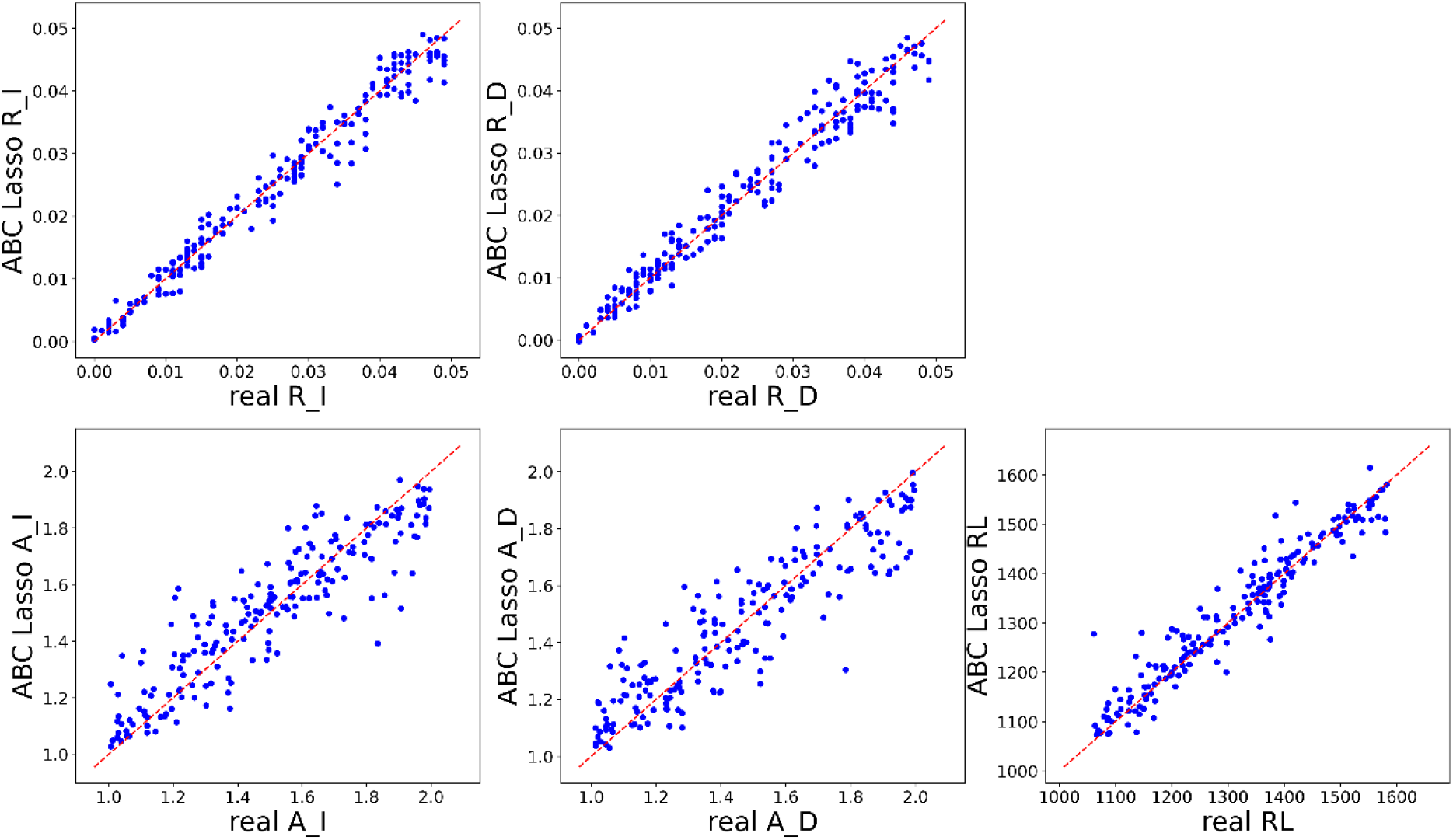
Model accuracy in simulations: inferred parameters (*R*_*I*, *R*_*D*, *A*_*I*,*A*_*D*, and *RL*) are correlated to the parameters used for simulation. Each point represents a single simulation inference for the corresponding parameter against the real value. Each graph is based on 200 independent simulations. The red dashed line is the identity *y* = *x* line. The obtained *R*^2^ values are 0.97, 0.96, 0.81, 0.83, and 0.92 for *R*_*I*, *R*_*D*, *A*_*I*, *A*_*D*, and *RL*, respectively.

### Feature importance

We use the terms “features” and “summary statistics” interchangeably. The impact of each summary statistic on the inference accuracy of SpartaABC was examined using simulations. SpartaABC computes 27 summary statistics for the input MSA and for each simulated MSA (table 2). The importance of each summary statistics for the inference accuracy of each of the five inferred parameters can be obtained from the Lasso regression coefficients. In Lasso regression, different features are assigned different weights, with greater penalty to high weights. This penalty prevents overfitting the data, practically reducing the number of features in each regression analysis (Tibshirani, 1996). Thus, the weight of each feature indicates its importance. A certain feature may be important for the inference of one parameter, but unimportant (zero weight) for another parameter. Moreover, some variability of feature importance is expected, to a certain degree, among datasets.

We computed a feature-importance score for each summary statistics for each of the five RIM parameters (supplementary fig. S2, Supplementary Material online). As expected, the most important feature for root length estimate was the alignment length. The second and third most important features were the lengths of the longest and the shortest sequences, respectively. For the insertion and deletion rate parameters, the most important summary statistics were those related to the number of gaps and alignment length. For example, the most important summary statistic for the *R*_*I* parameter was the total number of gaps in the alignment, while the most important summary statistic for the *R*_*D* parameter was the number of unique gaps (when counting unique gaps, those starting and ending at the same alignment position in more than one sequence, are only counted once). For the *A*_*I* parameter, which dictates the size distribution of newly inserted sequences, the most important summary statistic was the average gap size, and the second most important summary statistic was the number of gaps of length one. For the *A*_*D* parameter, which dictates the size distribution of new deletion events, the most important summary statistic was the number of gaps of length one in a single sequence. The feature importance analysis demonstrates the benefit of using multiple features for the accurate inference of parameters used in indel models. Furthermore, the regularization property of the Lasso regression methodology, may partially explain the improved accuracy of the Lasso-based inference compared to averaging over the posterior samples (supplementary table S1, Supplementary Material online).

### Model selection using neural networks

To compare the fit of different models, such as the SIM and RIM described above, to an empirical dataset, model selection procedures are needed. The most straightforward ABC model selection is to sample uniformly from the models (which is equivalent to assuming uniform prior over the models), pool all the simulations and select those that are closest to the empirical data, as defined by the distance threshold. The estimated posterior probability of each model is approximated by the relative frequency of retained simulations generated from each model (Pritchard et al., 1999). Alternative, more complex model selection procedures were previously developed (Bertorelle et al., 2010). However, it was previously shown that ABC model selection can be problematic under some scenarios (Roberta et al., 2011), and hence the performance of model selection procedures must be extensively tested using simulations. We compared two different model-selection schemes using simulations: the classic approach, and a neural-network-based classifier, which was recently developed in the field of population genetics (Mondal et al., 2019) and we implemented for indel-based models.

We first studied the power of both model selection procedures using simulations. To this end, for a given tree topology, we simulated 100 MSAs under various SIM parameters, as well as 100 MSAs under various RIM parameters. The classification confusion matrices for the dataset with seven sequences and mean sequence length of 1,384 amino acids, sampled from the EggNOG database, is shown in table 3. For this dataset, when the true model was SIM, both model-selection tests had similar high classification accuracy (above 97%). The differences between the two tests is shown when the dataset was simulated under RIM: the correct identification of the generating model was 82% for the neural-network based test and only 69% for the classic approach. These simulation results indicate that both model-selection tests slightly favor SIM over RIM, making the inference of RIM conservative. Of note, when simulating under RIM, the extent of the differences between the insertion and deletion parameters highly influenced the selected model. Indeed, the model-selection error was strongly dependent on the difference between *R*_*I* and *R*_*D* and to a lesser extent on the difference between *A*_*I* and *A*_*D* (fig. 3). The mean absolute difference between the *R*_*I* and *R*_*D* parameters for RIM simulations that were correctly classified as RIM was 0.018, while when the RIM simulations were misclassified as SIM it was 0.0034 (t-test P < 1e-16). The mean absolute difference between the *A*_*I* and *A*_*D* parameters for correctly classified RIM simulations was 0.36, while for RIM simulations misclassified as SIM it was 0.21 (t-test P < 0.02).

**Table 3.** Confusion matrices for model selection. Accuracy is computed based on 100 simulations, for each indel model. For example, out of 100 MSAs simulated under RIM, the classic model selection approach correctly identified 69 as RIM, while the neural network approach correctly identified 82.

| Classification method | Simulated model | Accuracy |
| --- | --- | --- |
| Classic | SIM | 0.98 |
| Classic | RIM | 0.69 |
| Neural network | SIM | 0.97 |
| Neural network | RIM | 0.82 |

**FIG. 2.**
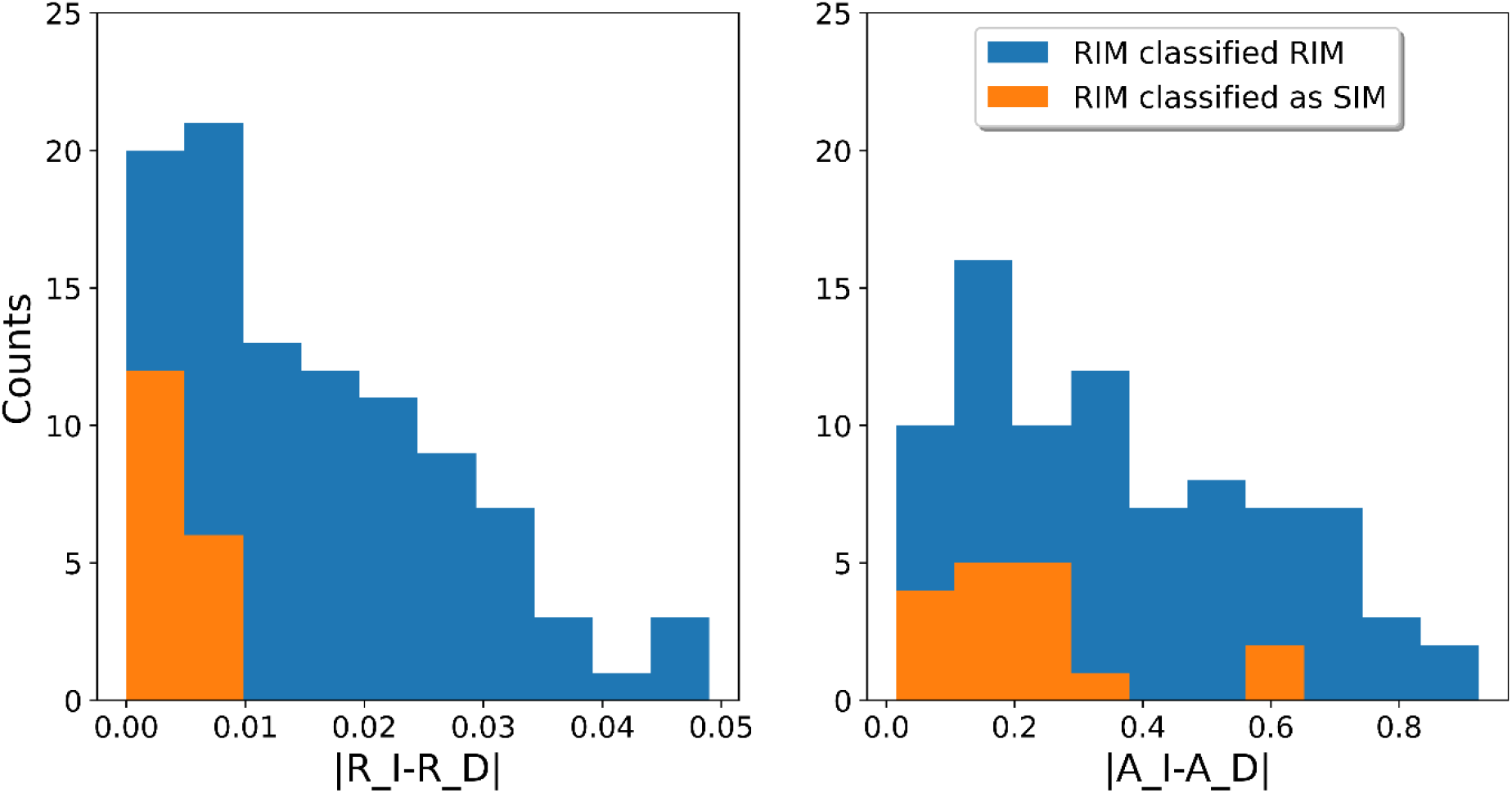
Misclassification rates depend on the similarity between insertion and deletion parameters. The errors depend on the absolute difference between *R*_*I* and *R*_*D* and the differences between *A*_*I* and *A*_*D*. All simulations were under the RIM model. In blue, simulations that were correctly classified as RIM and in orange, cases which were misclassified as SIM. Model selection was performed based on the neural-network classifier.

**FIG. 3.**
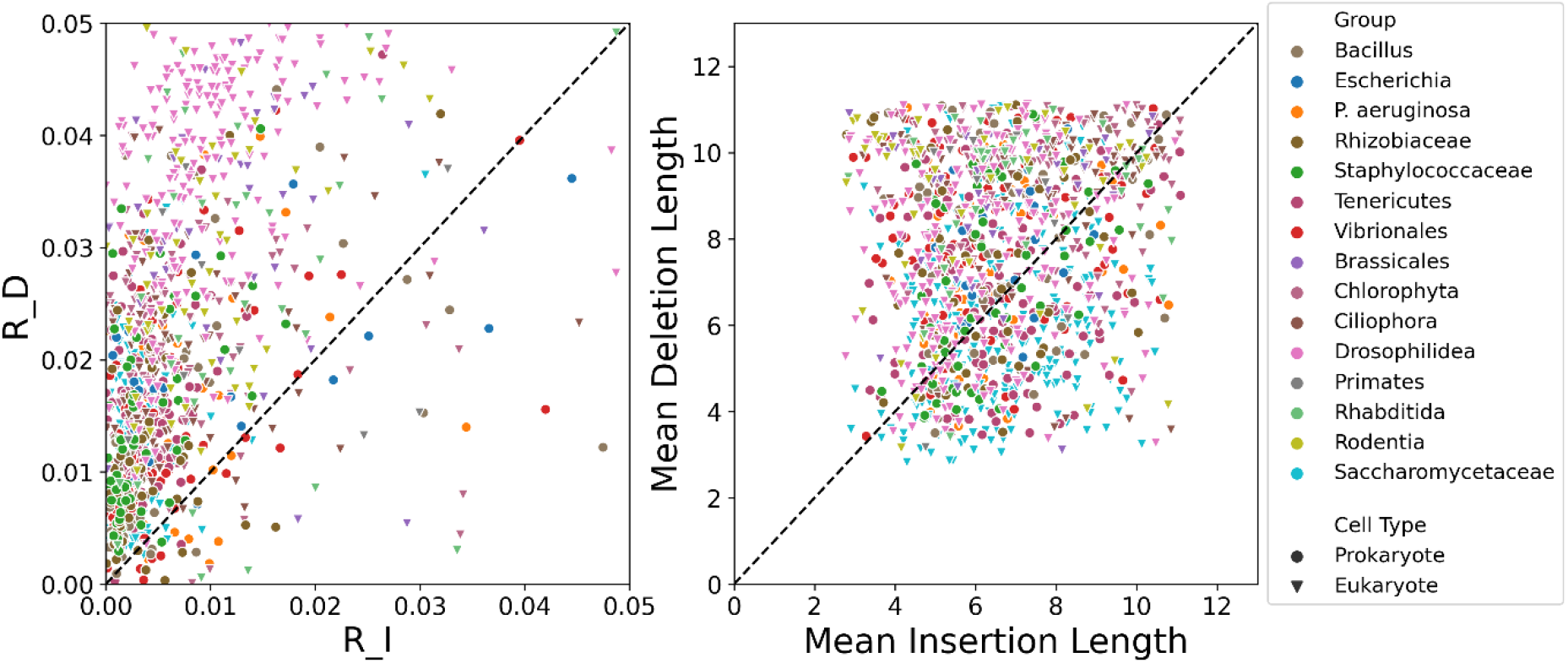
Deletion rates are mostly higher than insertion rates, while no significant trend is found for the length distribution. Left panel: a scatter plot of insertion rate (*R*_*I*) versus deletion rate (*R*_*D*). Right panel: a scatter plot of mean insertion length versus mean deletion rate. 1,232 datasets across 16 taxonomic groups for which the RIM model was selected are included in the analysis. The black dashed line is the identity line, *y* = *x*, in both panels.

We repeated this analysis for 12 additional datasets with various sequence lengths and total branch lengths. This yielded similar results regarding the classification errors: for the vast majority of the datasets the accuracy of the neutral-network and the classic procedures had similar accuracy in correctly identifying the SIM model. However, the neutral-network based model selection scheme had higher accuracy than the classic model in correctly identifying the RIM model (supplementary table S3, Supplementary Material online). Similar to the parameter inference accuracy, the model-selection test accuracy also depends on the total branch lengths (supplementary fig. S3, Supplementary Material online).

### Running times

The average running time for an empirical dataset was around 10 minutes on a single processor, including all simulations, extraction of features, and model selection between SIM and RIM. The running times for typical datasets are correlated to the total branch lengths of the examined phylogeny (*R*^2^ = 0.88, P < 2e-6): the running time in minutes is about five times the total branch length, when the branch lengths are measured in number of substitutions per site (supplementary fig. S4, Supplementary Material online).

### Empirical data analysis

We applied the model selection and inference algorithm on 2,649 biological datasets. These datasets included phylogenetic trees and protein MSAs of various phylogenetic groups, including bacteria, plants, insects, fungi, and mammals. Table 4 details the model-selection classifications for the various groups. Our method classified the model as RIM for 46% of the examined datasets. A large difference was observed between prokaryotic and eukaryotic organisms: for 62% of the eukaryotic datasets, RIM was selected over SIM, while for prokaryotic datasets, RIM was only selected in 33% of the cases. From the studied prokaryotes, the model was classified as RIM for the majority of the datasets only for Tenericutes. For most of the eukaryotic clades, the number of datasets for which SIM and RIM were selected was relatively equal; with the exceptions of Drosophilidea and Saccharomycetaceae for which the model was classified as RIM for the vast majority of the datasets. We note that it is likely that some of these differences do not reflect genuine differences between the two domains, but rather, differences in the data attributes (supplementary table S4, Supplementary Material online). For example, eukaryotic MSAs tend to be longer: the average length over all prokaryotic and eukaryotic datasets was 727.5 and 1101.7 amino acids, respectively (The average levels of tree divergence were relatively similar: the average sum of branch lengths over all prokaryotic and eukaryotic datasets was 8.2 and 7.2 amino-acid replacements per site, respectively).

**Table 4.** Model selection for various taxonomical groups. Pro/Euk: Prokaryotes/Eukaryotes.

| **Group** | **# SIM** | **# RIM** | **Pro/Euk** |
| --- | --- | --- | --- |
| *Bacillus* | 62 | 56 | Pro |
| *Escherichia* | 103 | 20 | Pro |
| *P. aeruginosa* | 178 | 53 | Pro |
| *Rhizobiaceae* | 132 | 54 | Pro |
| *Staphylococcaceae* | 227 | 60 | Pro |
| Tenericutes | 83 | 123 | Pro |
| Vibrionales | 145 | 88 | Pro |
| Brassicales | 53 | 40 | Euk |
| Chlorophyta | 93 | 100 | Euk |
| Ciliophora | 87 | 82 | Euk |
| Drosophilidea | 39 | 202 | Euk |
| Primates | 17 | 9 | Euk |
| Rhabditida | 93 | 36 | Euk |
| Rodentia | 45 | 48 | Euk |
| Saccharomycetaceae | 60 | 261 | Euk |

The mean values of the various model parameters, per taxonomic group, for the RIM and SIM selected datasets are shown in table 5a and table 5b, respectively. For brevity, the mean insertion and deletion lengths are given instead of the power law parameters. *R*_*D* was higher than *R*_*I* for all examined taxonomic groups (table 5a). *Staphylococcaceae*, Tenericutes, Drosophilidea, and Saccharomycetaceae were found to have the highest deletion-insertion ratio, i.e., *R*_*D*⁄*R*_*I*. As expected, most of the datasets for these four groups were classified as RIM.

**Table 5.** Model parameters across various taxonomical groups for datasets classified as RIM (A) and SIM (B).

| Group | *RL* | *R_I* | *R_D* | Mean<br>insertion<br>length | Mean<br>deletion<br>length |
| --- | --- | --- | --- | --- | --- |
| *Bacillus* | 808.6 | 0.0071 | 0.0146 | 6.83 | 8.24 |
| *Escherichia* | 491.8 | 0.0111 | 0.0192 | 6.67 | 7.54 |
| *P. aeruginosa* | 803.1 | 0.0059 | 0.0154 | 6.31 | 7.22 |
| *Rhizobiaceae* | 844.5 | 0.0043 | 0.0116 | 6.30 | 7.53 |
| *Staphylococcaceae* | 645.7 | 0.0038 | 0.0139 | 6.39 | 7.33 |
| Tenericutes | 914.5 | 0.0039 | 0.0154 | 6.59 | 6.97 |
| Vibrionales | 797.4 | 0.0057 | 0.0133 | 6.33 | 8.28 |
| Brassicales | 1406.8 | 0.0120 | 0.0324 | 6.57 | 8.83 |
| Chlorophyta | 1043.5 | 0.0073 | 0.0206 | 7.84 | 9.36 |
| Ciliophora | 1022.4 | 0.0078 | 0.0202 | 7.28 | 8.79 |
| Drosophilidea | 1745.5 | 0.0099 | 0.0361 | 6.13 | 7.64 |
| Primates | 1598.1 | 0.0147 | 0.0240 | 6.45 | 8.56 |
| Rhabditida | 1053.0 | 0.0128 | 0.0259 | 7.33 | 9.10 |
| Rodentia | 1084.1 | 0.0097 | 0.0284 | 6.10 | 8.72 |
| Saccharomycetaceae | 1193.7 | 0.0027 | 0.0110 | 6.56 | 6.11 |

B.
| Group | *RL* | *R_ID* | Mean<br>indel<br>length |
| --- | --- | --- | --- |
| *Bacillus* | 667.3 | 0.0266 | 9.14 |
| *Escherichia* | 443.3 | 0.0190 | 6.56 |
| *P. aeruginosa* | 810.1 | 0.0154 | 7.20 |
| *Rhizobiaceae* | 873.1 | 0.0118 | 7.34 |
| *Staphylococcaceae* | 572.0 | 0.0094 | 6.37 |
| Tenericutes | 673.9 | 0.0178 | 7.76 |
| Vibrionales | 677.7 | 0.0156 | 7.67 |
| Brassicales | 1025.8 | 0.0514 | 8.88 |
| Chlorophyta | 717.7 | 0.0352 | 9.63 |
| Ciliophora | 618.5 | 0.0362 | 9.75 |
| Drosophilidea | 1198.3 | 0.0532 | 8.25 |
| Primates | 970.0 | 0.0636 | 8.48 |
| Rhabditida | 679.4 | 0.0540 | 9.24 |
| Rodentia | 717.6 | 0.0500 | 8.93 |
| Saccharomycetaceae | 1181.9 | 0.0088 | 6.35 |

Figure 3 shows scatter plots of *R*_*D* vs. *R*_*I* and mean deletion length vs. mean insertion length for the datasets classified as RIM. In most of these datasets (1,156 out of 1,232), the deletion rate was higher than the insertion rate. The mean deletion length tended to be higher than the insertion lengths, however, this trend is quite insubstantial.

Sung et al. (2016) showed that the effective population size is highly correlated to the indel rate. Hence, we aimed to test if such dependence is observed when using our probabilistic indel-based models. For our analysis, we selected taxonomic groups from the EggNOG database (Huerta-Cepas et al., 2019) that are as similar as possible to the ones in the Sung et al. (2016) study. We note that in the current analysis, indel rates were inferred for proteins rather than for neutrally evolving regions. Another difference is that the inferred rates, *R*_*I* and *R*_*D*, are relative to the protein substitution rate. Despite these differences, we observed a high correlation (*R*^2^ = 0.61) between the total indel rate (*R*_*I* + *R*_*D*) and the effective population size (fig. 4, left panel).

**FIG. 4.**
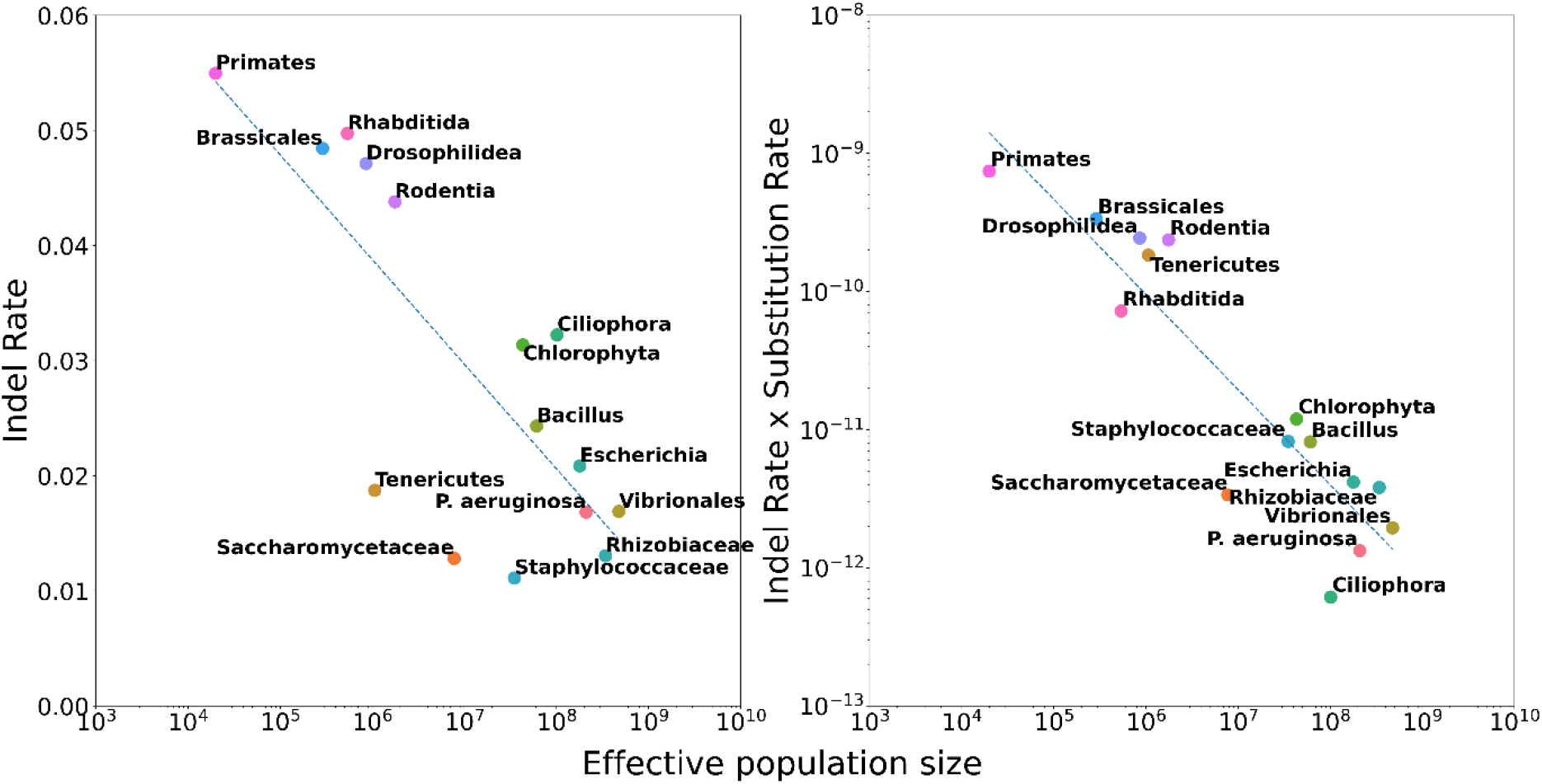
Indel rate as a function of effective population size. Effective population size (*EFP*) vs. Indel-to-substitution rate (left panel) and effective population size vs. Indel-to-substitution rate × Substitution rate (right panel). For each taxonomic group, the total indel rate reflects the average over all datasets for this group. The substitution rates and effective population sizes are taken from Sung et al. (2016). The dotted regression lines are: (a) Indel-to-substitution rate = −0.0091 × *log*_10_*EFP* + 0.093, (*R*^2^ = 0.61, P = 6E-4); (b) *log*_10_[Indel-to-substitution rate × Substitution rate] = −0.69 × *log*_10_*EFP* − 5.90, (*R*^2^ = 0.83, P = 3E-6).

In a different study, Sung et al. (2012) showed a correlation between the substitution rate and the effective population size. In addition, a high correlation was inferred between the indel and substitution rates (Sung et al., 2016). In our probabilistic models, the indel rates are computed relative to the substitution rate. To evaluate the correlation between effective population sizes and indel rates, without normalizing the substitution rate, we repeated the correlation analysis, this time contrasting the effective population size against the total indel rate multiplied by the substitution rate (fig. 4, right panel). Substitution rates were taken from Sung et al. (2012). We note that the substitution rates reported in Sung et al. (2012) are for nucleotide sequences, and the substitution rates in our analysis are in number of amino-acid replacements per site, yet this should not affect the correlation analysis as long as these two values are proportional. The observed correlation in this case was even higher, *R*^2^ = 0.83, compared to the case when indel rate is normalized to the substitution rate.

## Discussion

In this work we developed an indel model that accounts for differences between insertion and deletion evolutionary dynamics. Furthermore, we developed an ABC inference scheme to estimate model parameters as well as a model-selection test, using a neural-network classifier, that is able to determine which model (RIM or SIM) better fits a given empirical dataset. In our simulations, both the model selection and the inference steps were shown to be accurate. Applying the developed inference scheme on a variety of empirical datasets allowed us to gain further insights on indel dynamics. First, for 46% of the examined datasets, the inferred insertions and deletions rates and length distributions were different. Interestingly, this fraction was larger for eukaryotic than prokaryotic organisms. Second, the deletion rate is typically larger than the insertion rate (and to a much lesser extent, the deletion length is larger than the insertion length). Third, we compared the indel rates of various organisms to the effective population size and showed that they were negatively correlated.

The RIM model established in this study is more elaborate than our previous model (SIM) that assumed equal attributes of insertions and deletions (Karin et al., 2017). It was previously shown for both prokaryotes and eukaryotes that there is a deletion bias on sites which are assumed to evolve under neutral selection (Kuo & Ochman, 2009; Mira et al., 2001; Ophir & Graur, 1997; Petrov et al., 1996; Van Passel et al., 2007; Zhang & Gerstein, 2003). Here, we showed that for a large variety of phylogenetic clades there is a deletion bias also for coding protein sequences, which generally evolve under strong purifying selection and that this bias is mainly due to high deletion rates, rather than due to longer deletion events. Of note, in our models the estimated deletion rates are normalized by the substitution rate. Often, when comparing two organisms, one is inferred to have both higher deletion to substitution rate and higher substitution rate. Together these two factors result in markedly different deletion dynamics, which may have impact on genome sizes (Petrov et al., 2000).

The indel models developed here have several limitations and there is still much room for more realistic modeling extensions. For example, both SIM and RIM assume that the indel parameters are uniform across the whole input sequence. However, it was shown that the composition of amino acids in and around indels is significantly different from their composition across the entire sequence length, with enrichment for amino acids ADQEGPS and depletion of FMILYVWC (Chang & Benner, 2004). Other studies also demonstrated that indel rates depend on the amino acid context (De La Chaux, Messer, & Arndt, 2007; Kvikstad, Chiaromonte, & Makova, 2009; Kvikstad & Duret, 2014; Messer & Arndt, 2007; Tanay & Siggia, 2008). Ideally, empirical context-dependent indel models should be developed, in which the rate of insertions and the rate of deletions should each depend on the amino acid composition surrounding the indel site. Such models are expected to have a large number of free parameters, and thus resemble empirical amino acid replacement matrices such as JTT (Jones, Taylor, & Thornton, 1992), WAG (Whelan & Goldman, 2001), and LG (Le & Gascuel, 2008). Accurate inference of the model parameters would require simultaneous analysis of a large amount of data, e.g., the entire set of mammalian MSAs. High quality genomic data become increasingly available, and provide fertile ground for the development of such models. Of note, the ABC inference scheme described above relies on efficient simulators. To accelerate parameter inference we implemented a simulator that generates indel events only and does not include substitution events. The above context dependent models will necessitate simulating indel and substitution events simultaneously.

Another direction for future advance is to develop indel models that account for structural features of protein-coding genes. It is expected that different structural attributes do not share the same indel dynamics. For example, it was recently shown that for enhanced green fluorescent protein (eGFP), the packing density of a residue, as measured by the weighted contact number (Lin et al., 2008), considerably affects the probability that a single-residue deletion disrupts the protein’s function (Jackson et al., 2017). Relative surface accessibility and secondary structure attributes were also found to affect this probability. Future indel models can explicitly account for such factors and should prove particularly useful, providing that the secondary or tertiary structures of a protein are available or can be accurately predicted.

Most commonly used alignment algorithms maximize a specific score and do not explicitly assume a stochastic Markov process. Recently, advances have been made in the development of statistical alignment methods, allowing simultaneous model parameter inference and alignment (Suchard & Redelings, 2006; Novák et al., 2008; Bradley et al., 2009; Nute et al., 2019). Miklós et al. (2004) and Levy-Karin et al. (2019) have developed the long-indel model, in which both indels and substitutions evolve along a phylogenetic tree assuming a joint continuous time Markov process. The rate of indels in this model depends on the indel length. Such a model was shown to better fit empirical data compared to previous models such as TKF91 (Thorne et al., 1991). However, it requires extensive computational time and is currently limited to pairwise sequences. This model also assumes that deletion and insertion events have the same dynamics. The results of this study corroborate previous studies showing that for a large number of empirical datasets, insertion and deletion events are characterized by different evolutionary dynamics. Such considerations should be included in future statistical alignment methodologies.

The applicability of our methodology was demonstrated here by corroborating a previously observed correlation between effective population sizes and indel rates. Of note, in this study the indel rates are computationally inferred from a known set of protein coding genes, while Sung et al. (2016) inferred it using whole genome sequencing of mutation-accumulation experiments. An additional possible application of our methodology would be to characterize how indel dynamics vary among different proteins and protein domains, e.g., it was previously suggested that ancient protein domains (Wolf et al., 2007) and highly conserved proteins (Ajawatanawong & Baldauf, 2013) have a bias towards insertions and that essential proteins in bacteria and yeast experience more indel events than non-essential proteins (Chan et al., 2007). Finally, our methodology can be applied at the DNA level to quantify and test hypotheses regarding indel dynamics in introns, promotors, enhancers, regions with high or low recombination rates, etc., and should provide statistical-sound means to compare indel dynamics among genes and genomes across the tree of life.

## Materials and methods

### Source code and implementation details

The implementation of the algorithm presented here is called SpartaABC. It is implemented in C++ and Python. The source code is freely available in https://github.com/gilloe/SpartaABC. The following Python packages were used for machine learning: scikit-learn and keras. The SIM and RIM models, including the model selection schemes were added to the SPARTA ABC webserver: https://spartaabc.tau.ac.il/ (Ashkenazy et al., 2017).

### Empirical datasets

The data used to generate figures 1 and 2 are based on the tree and MSA available from EggNOG (Huerta-Cepas et al., 2019) entry Drosophilidea ENOG410HUGV. Supplementary table S1 is based on the analysis of 12 additional EggNOG datasets (supplementary table S1, Supplementary Material online). The data used for the feature importance analyses are based on six EggNOG datasets (supplementary fig. S2, Supplementary Material online). The data used to generate table 3 and figure 2 are based on the tree and MSA EggNOG entry Drosophilidea ENOG410HUGV. The biological datasets, i.e. the empirical phylogenetic trees and MSAs, were also downloaded from EggNOG. Due to computational limitations, and in order that each taxonomic group will contain similar number of datasets, inclusion criteria were applied. Specifically, we determined a minimal MSA length and a minimal number of species for each taxonomic group (supplementary table S4, Supplementary Material online). In addition, we filtered datasets in which the total branch lengths of the tree was smaller than 1 (supplementary fig. S1, Supplementary Material online). We also filtered out datasets in which the inference results fell outside the prior.

### A neural-network classifier for model selection

A neural-network classifier, similar to Mondal et al. (2019), was used in order to determine whether RIM, in which the rate and length distribution are allowed to differ between insertions and deletions, is more suitable for a particular dataset than SIM, in which the insertion and deletion rates, as well as insertions and deletions length distributions, are equal (Karin et al., 2017). The classifier was trained for a specific phylogenetic tree and MSA using 100,000 simulations from each model (the same simulations were used for parameter inference and model selection). The normalized summary statistics were the input for the classifier and the output was the probability per model, such that the model with the highest probability was selected. The architecture of this classifier is shown in supplementary fig. S5, Supplementary Material online.

### Model-selection accuracy

For assessing the model-selection accuracy for a specific phylogenetic tree and MSA (note that the MSA is used only for setting the *RL* prior), we simulated 200 datasets per model (400 in total) by randomly drawing combinations of parameters from the prior distributions of these models. Then, we used the model-selection scheme to infer the parameters of the simulated datasets and calculated the confusion matrix using the inference results and the known model from which the samples were drawn.

## Acknowledgment

BSF [2015247 to RAC, TP]; ISF [802/16 to TP]; GL, DR, OA, AM, and DA were supported in part by a fellowship from the Edmond J. Safra Center for Bioinformatics at Tel Aviv University. OA was partly supported by the Dalia and Eli Hurvits foundation.

