## Supplementary Material for "A probabilistic model for indel evolution: differentiating insertions from deletions"

### Supplementary Information

Gil Loewenthal[1], Dana Rapoport[1], Oren Avram[1], Asher Moshe[1], Alon Itzkovitch[1], Omer Israeli[1], Dana Azouri[1,2], Reed A. Cartwright[3,4], Itay Mayrose[2], and Tal Pupko[1,†]

[1] The Shmunis School of Biomedicine and Cancer Research, George S. Wise Faculty of Life Sciences, Tel Aviv University, Tel Aviv 69978, Israel.

[2] School of Plant Sciences and Food Security, George S. Wise Faculty of Life Sciences, Tel Aviv University, Tel Aviv 69978, Israel.

[3] The Biodesign Institute, Arizona State University, Tempe, Arizona, USA.

[4] School of Life Sciences, Arizona State University, Tempe, Arizona, USA.

† To whom correspondence should be addressed:

**Table S1.** A comparison between various ABC inference schemes: Mean, Ridge, Lasso.

| Parameter | Mean – $R^2$ | Ridge - $R^2$ | Lasso - $R^2$ |
|---|---|---|---|
| RL | 0.86 | 0.93 | 0.92 |
| IR | 0.95 | 0.97 | 0.97 |
| DR | 0.95 | 0.96 | 0.96 |
| AI | 0.77 | 0.80 | 0.81 |
| AD | 0.79 | 0.83 | 0.83 |

**Table S2.** Dataset comparison. Shown are the dataset topology parameters with different simulation accuracy metrics with Lasso regression. The slope was derived from regression by setting the intercept to zero.

| Group | EggNOG accession id | Sum of branch lengths [aa substitutions] | MSA average length [aa] | Number of sequences | Parameter | $R^2$ | Slope |
|---|---|---|---|---|---|---|---|
| Artropoda | ENOG41032B6 | 16.91 | 1,597 | 22 | RL | 0.98 | 1.01 |
| | | | | | R_I | 0.98 | 0.98 |
| | | | | | R_D | 0.97 | 1.00 |
| | | | | | A_I | 0.90 | 1.00 |
| | | | | | A_D | 0.85 | 0.99 |
| Brassicales | ENOG410C1CI | 0.78 | 735 | 5 | RL | 0.92 | 1.00 |
| | | | | | R_I | 0.80 | 0.94 |
| | | | | | R_D | 0.70 | 0.90 |
| | | | | | A_I | 0.37 | 0.97 |
| | | | | | A_D | 0.48 | 0.99 |
| Chlorobi | ENOG410CZAT | 2.31 | 564 | 11 | RL | 0.62 | 0.99 |
| | | | | | R_I | 0.86 | 0.95 |
| | | | | | R_D | 0.85 | 0.95 |
| | | | | | A_I | 0.55 | 0.97 |
| | | | | | A_D | 0.51 | 0.99 |
| Chlorobi | ENOG410CZH7 | 5.66 | 626 | 12 | RL | 0.60 | 1.00 |
| | | | | | R_I | 0.95 | 0.99 |
| | | | | | R_D | 0.93 | 0.99 |
| | | | | | A_I | 0.68 | 0.99 |
| | | | | | A_D | 0.67 | 0.99 |
| Cyanobacteria | ENOG410FNZW | 9.47 | 1,216 | 10 | RL | 0.96 | 1.00 |
| | | | | | R_I | 0.98 | 0.99 |
| | | | | | R_D | 0.97 | 0.99 |
| | | | | | A_I | 0.83 | 0.99 |
| | | | | | A_D | 0.78 | 1.00 |
| Drosophilidea | ENOG410HUGV | 4.66 | 1,384 | 7 | RL | 0.92 | 1.01 |
| | | | | | R_I | 0.97 | 0.99 |
| | | | | | R_D | 0.96 | 0.99 |
| | | | | | A_I | 0.81 | 1.00 |

| | | | | | | | |
|---|---|---|---|---|---|---|---|
| | | | | | A_D | 0.83 | 0.99 |
| Epsilonproteobacteria | ENOG410I2VE | 13.32 | 727 | 41 | RL | 0.90 | 1.00 |
| | | | | | R_I | 0.98 | 0.99 |
| | | | | | R_D | 0.97 | 0.99 |
| | | | | | A_I | 0.86 | 0.99 |
| | | | | | A_D | 0.80 | 0.99 |
| Fish | ENOG410N476 | 4.53 | 1,278 | 8 | RL | 0.98 | 0.99 |
| | | | | | R_I | 0.89 | 1.00 |
| | | | | | R_D | 0.91 | 0.97 |
| | | | | | A_I | 0.65 | 0.98 |
| | | | | | A_D | 0.72 | 0.99 |
| Fungi | ENOG410PK11 | 19.72 | 1,194 | 20 | RL | 0.93 | 1.00 |
| | | | | | R_I | 0.97 | 1.00 |
| | | | | | R_D | 0.97 | 0.99 |
| | | | | | A_I | 0.89 | 1.00 |
| | | | | | A_D | 0.82 | 0.99 |
| Primates | ENOG411668K | 0.25 | 1,709 | 6 | RL | 0.99 | 1.00 |
| | | | | | R_I | 0.66 | 0.92 |
| | | | | | R_D | 0.62 | 0.92 |
| | | | | | A_I | 0.32 | 0.98 |
| | | | | | A_D | 0.32 | 0.97 |
| Primates | ENOG4116ES6 | 0.5 | 1,855 | 7 | RL | 0.96 | 1.00 |
| | | | | | R_I | 0.69 | 0.93 |
| | | | | | R_D | 0.69 | 0.90 |
| | | | | | A_I | 0.36 | 0.97 |
| | | | | | A_D | 0.32 | 0.97 |
| Rodents | ENOG4118Z5J | 1.74 | 1,341 | 5 | RL | 0.95 | 1.00 |
| | | | | | R_I | 0.92 | 0.97 |
| | | | | | R_D | 0.93 | 0.97 |
| | | | | | A_I | 0.68 | 0.99 |
| | | | | | A_D | 0.66 | 1.01 |
| Rodents | ENOG41190IG | 0.23 | 1,299 | 5 | RL | 1.00 | 1.00 |
| | | | | | R_I | 0.69 | 0.90 |
| | | | | | R_D | 0.71 | 0.88 |
| | | | | | A_I | 0.30 | 0.97 |
| | | | | | A_D | 0.18 | 0.99 |

**Table S3.** The confusion matrices of the classic and neural-network (NN) classifiers. The accuracy (number of cases for which the model selection classifier correctly identified the generative model) analysis is shown for 13 empirical datasets. For each dataset, 200 simulations were generated (100 SIM and 100 RIM).

| Group | Eggnog accession id | SIM Accuracy Classic | SIM Accuracy NN | RIM Accuracy Classic | RIM Accuracy NN |
|---|---|---|---|---|---|
| Artropoda | ENOG41032B6 | 100 | 99 | 73 | 89 |
| Brassicales | ENOG410C1CI | 79 | 79 | 58 | 65 |
| Chlorobi | ENOG410CZAT | 90 | 86 | 55 | 69 |
| Chlorobi | ENOG410CZH7 | 93 | 94 | 59 | 64 |
| Cyanobacteria | ENOG410FNZW | 99 | 96 | 66 | 80 |
| Drosophilidea | ENOG410HUGV | 98 | 97 | 69 | 82 |
| Epsilonproteobacteria | ENOG410I2VE | 97 | 93 | 73 | 90 |
| Fish | ENOG410N476 | 94 | 92 | 63 | 82 |
| Fungi | ENOG410PK11 | 99 | 98 | 65 | 89 |
| Primates | ENOG411668K | 74 | 72 | 60 | 58 |
| Primates | ENOG4116ES6 | 59 | 71 | 63 | 53 |
| Rodents | ENOG4118Z5J | 96 | 94 | 60 | 74 |
| Rodents | ENOG41190IG | 82 | 85 | 52 | 45 |

**Table S4**. Inclusion thresholds per taxonomic group

| Group | MSA length | Number of species |
|---|---|---|
| *Bacillus* | 800 | 8 |
| *Escherichia* | 300 | 5 |
| *P. aeruginosa* | 700 | 4 |
| *Rhizobiaceae* | 800 | 8 |
| *Staphylococcaceae* | 500 | 5 |
| Tenericutes | 700 | 4 |
| Vibrionales | 700 | 7 |
| Brassicales | 1,500 | 7 |
| Chlorophyta | 1,000 | 10 |
| Ciliophora | 800 | 8 |
| Drosophilidea | 1,500 | 10 |
| Primates | 1,500 | 10 |
| Rhabditida | 1,000 | 10 |
| Rodentia | 1,000 | 15 |
| Saccharomycetaceae | 1,000 | 15 |

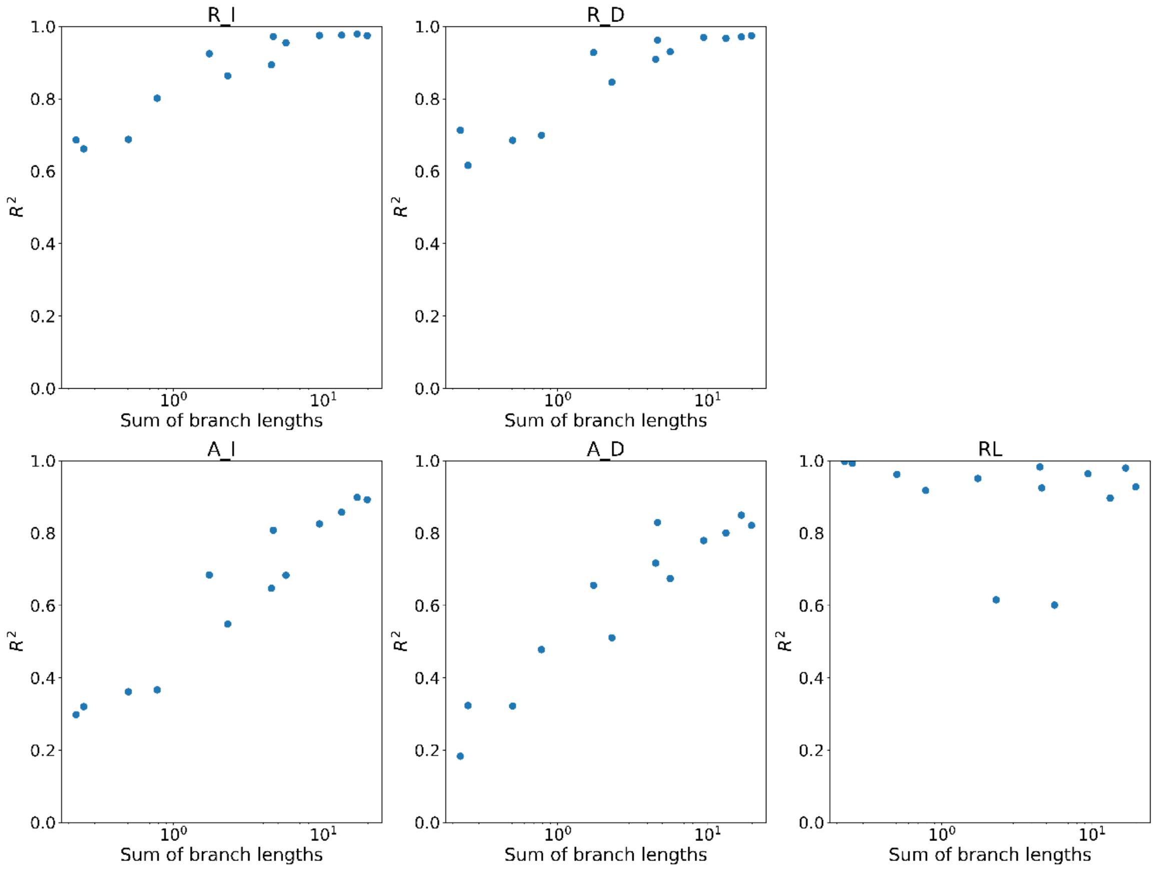


**Figure S1.** Scatter plots of the $R^2$ of the simulation set (from table S2) versus the sum of branch lengths for the inferred parameters (R_I, R_D, A_I, A_D, RL).

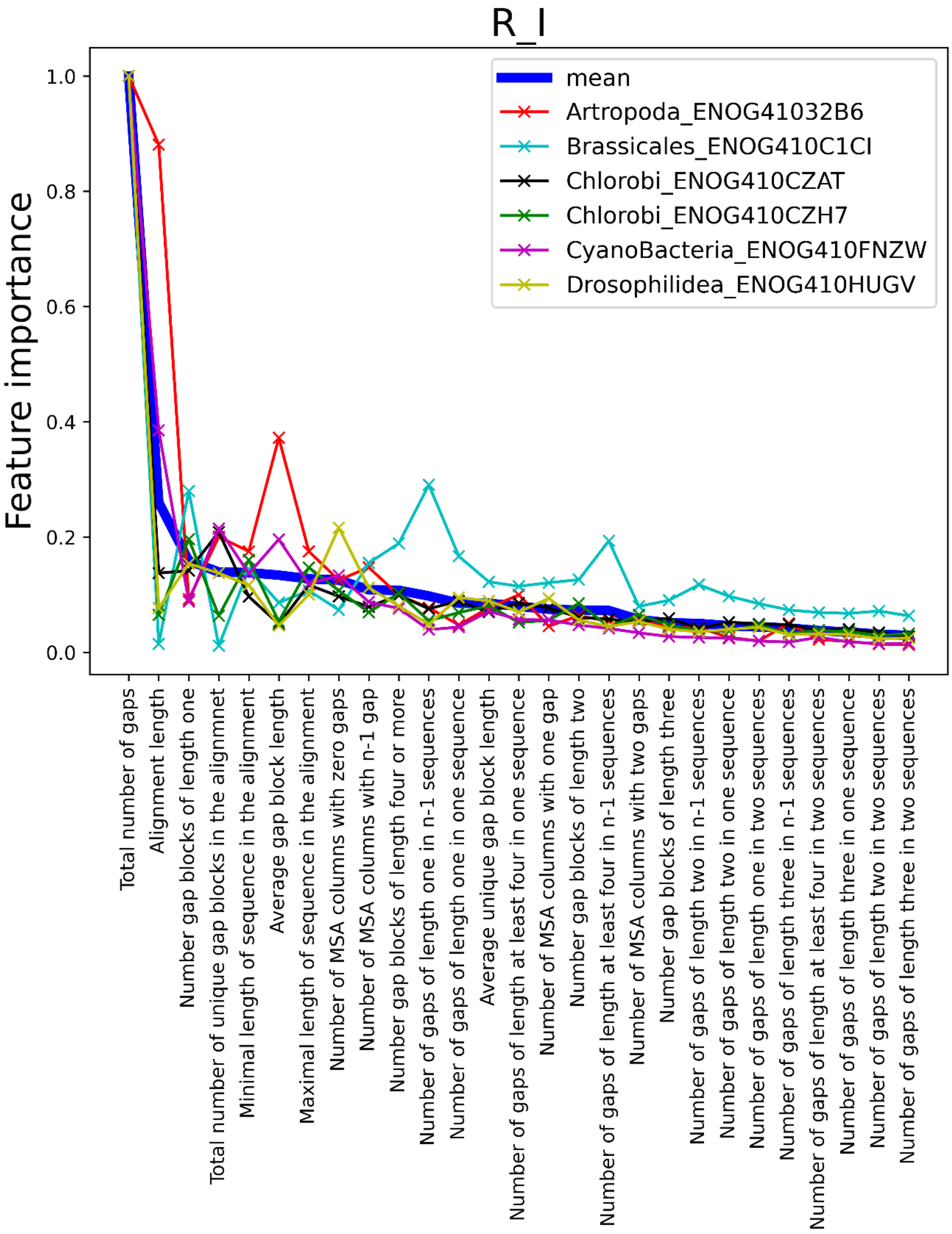

R_I
Feature importance
1.0
0.8
0.6
0.4
0.2
0.0
mean
Artropoda_ENOG41032B6
Brassicales_ENOG410C1CI
Chlorobi_ENOG410CZAT
Chlorobi_ENOG410CZH7
CyanoBacteria_ENOG410FNZW
Drosophilidea_ENOG410HUGV
Total number of gaps
Alignment length
Number gap blocks of length one
Total number of unique gap blocks in the alignmnet
Minimal length of sequence in the alignment
Average gap block length
Maximal length of sequence in the alignment
Number of MSA columns with zero gaps
Number of MSA columns with n-1 gap
Number gap blocks of length four or more
Number of gaps of length one in n-1 sequences
Number of gaps of length one in one sequence
Average unique gap block length
Number of gaps of length at least four in one sequence
Number of MSA columns with one gap
Number gap blocks of length two
Number of gaps of length at least four in n-1 sequences
Number of MSA columns with two gaps
Number gap blocks of length three
Number of gaps of length two in n-1 sequences
Number of gaps of length two in one sequence
Number of gaps of length one in two sequences
Number of gaps of length three in n-1 sequences
Number of gaps of length at least four in two sequences
Number of gaps of length three in one sequence
Number of gaps of length two in two sequences
Number of gaps of length three in two sequences

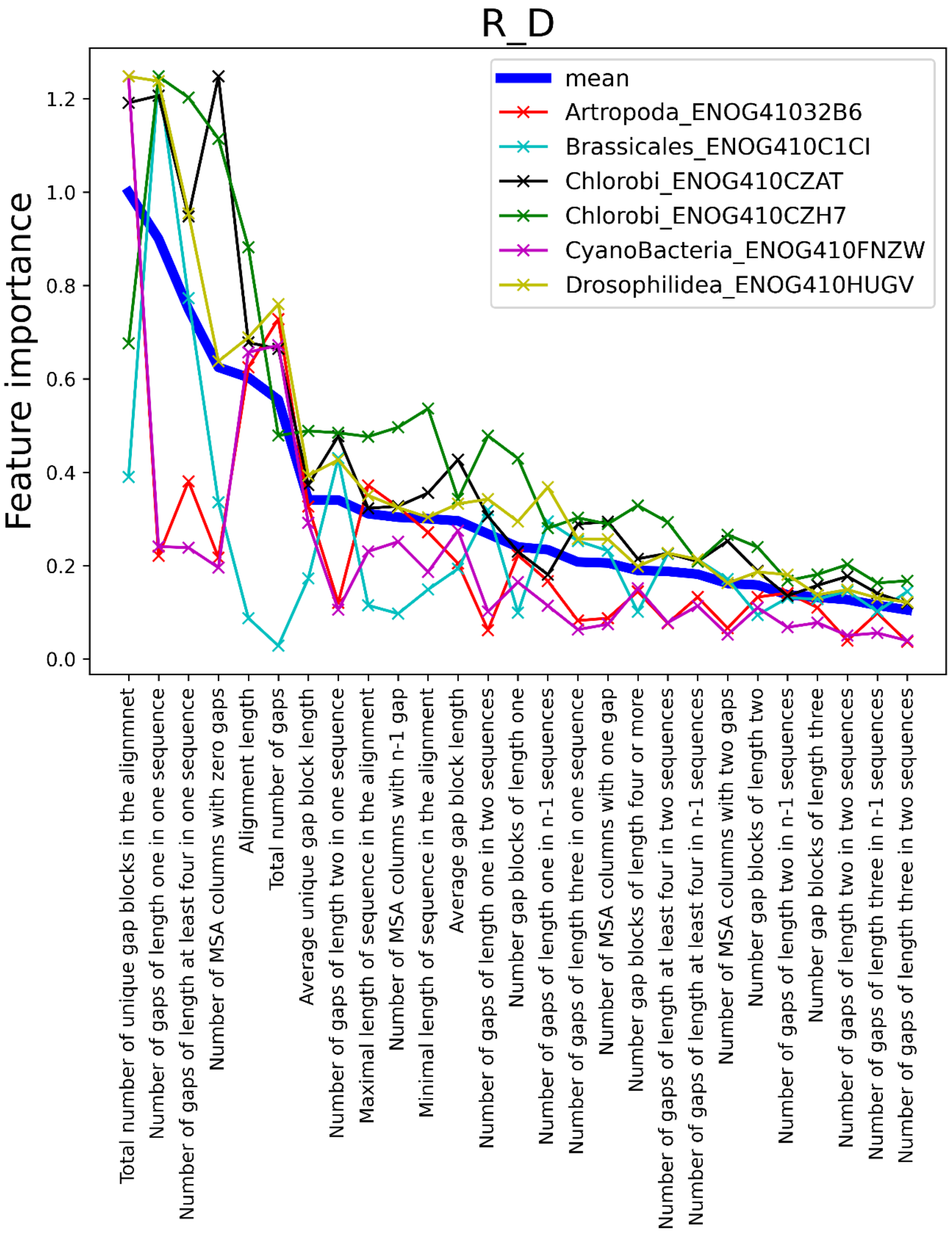
R_D
Feature importance
mean
Artropoda_ENOG41032B6
Brassicales_ENOG410C1CI
Chlorobi_ENOG410CZAT
Chlorobi_ENOG410CZH7
CyanoBacteria_ENOG410FNZW
Drosophilidea_ENOG410HUGV
0.0
0.2
0.4
0.6
0.8
1.0
1.2
Total number of unique gap blocks in the alignmnet
Number of gaps of length one in one sequence
Number of gaps of length at least four in one sequence
Number of MSA columns with zero gaps
Alignment length
Total number of gaps
Average unique gap block length
Number of gaps of length two in one sequence
Maximal length of sequence in the alignment
Number of MSA columns with n-1 gap
Minimal length of sequence in the alignment
Average gap block length
Number of gaps of length one in two sequences
Number gap blocks of length one
Number of gaps of length one in n-1 sequences
Number of gaps of length three in one sequence
Number of MSA columns with one gap
Number gap blocks of length four or more
Number of gaps of length at least four in two sequences
Number of gaps of length at least four in n-1 sequences
Number of MSA columns with two gaps
Number gap blocks of length two
Number of gaps of length two in n-1 sequences
Number gap blocks of length three
Number of gaps of length two in two sequences
Number of gaps of length three in n-1 sequences
Number of gaps of length three in two sequences

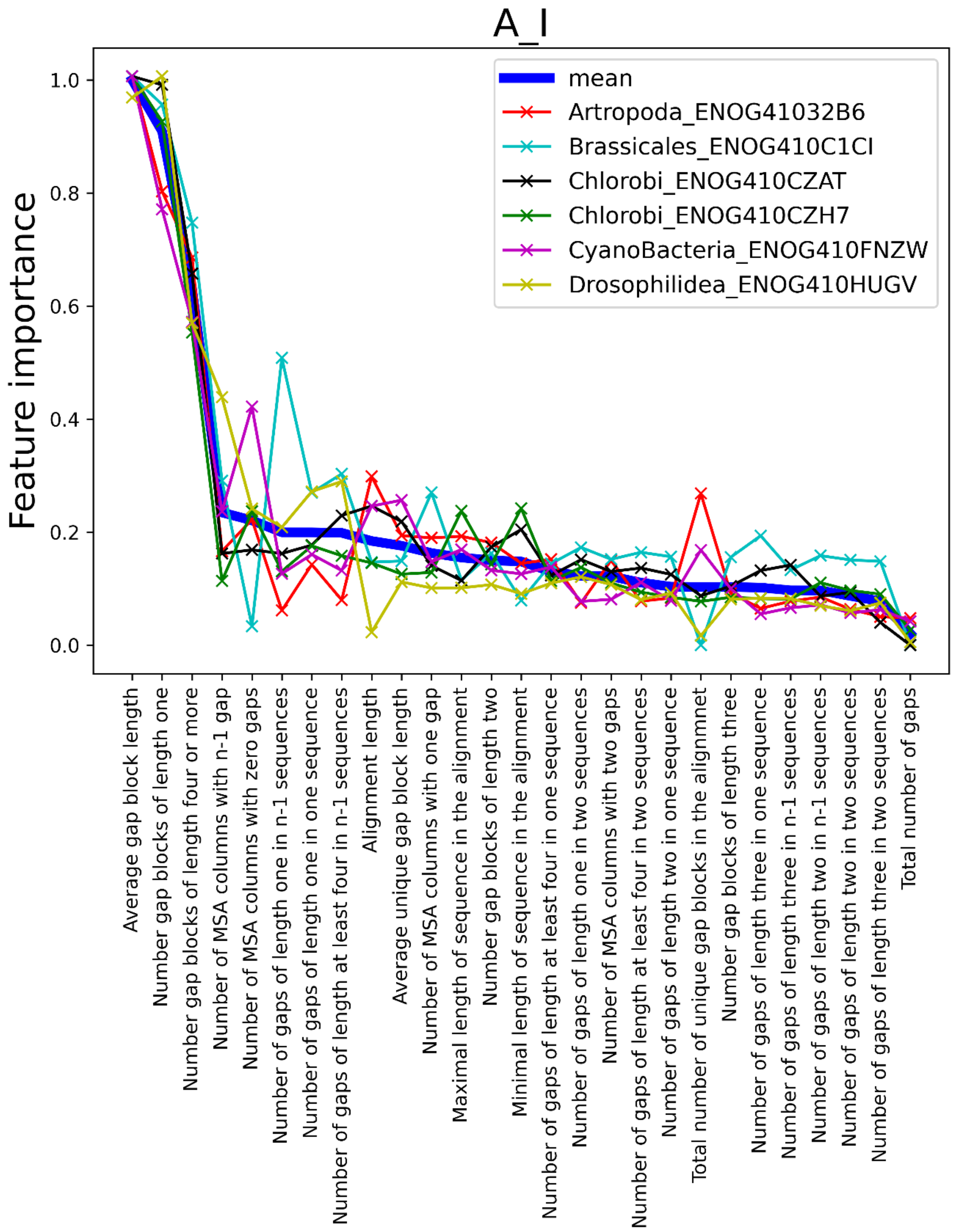

A_I
Feature importance
1.0
0.8
0.6
0.4
0.2
0.0
mean
Artropoda_ENOG41032B6
Brassicales_ENOG410C1CI
Chlorobi_ENOG410CZAT
Chlorobi_ENOG410CZH7
CyanoBacteria_ENOG410FNZW
Drosophilidea_ENOG410HUGV
Average gap block length
Number gap blocks of length one
Number gap blocks of length four or more
Number of MSA columns with n-1 gap
Number of MSA columns with zero gaps
Number of gaps of length one in n-1 sequences
Number of gaps of length one in one sequence
Number of gaps of length at least four in n-1 sequences
Alignment length
Average unique gap block length
Number of MSA columns with one gap
Maximal length of sequence in the alignment
Number gap blocks of length two
Minimal length of sequence in the alignment
Number of gaps of length at least four in one sequence
Number of gaps of length one in two sequences
Number of MSA columns with two gaps
Number of gaps of length at least four in two sequences
Number of gaps of length two in one sequence
Total number of unique gap blocks in the alignmnet
Number gap blocks of length three
Number of gaps of length three in one sequence
Number of gaps of length three in n-1 sequences
Number of gaps of length two in n-1 sequences
Number of gaps of length two in two sequences
Number of gaps of length three in two sequences
Total number of gaps

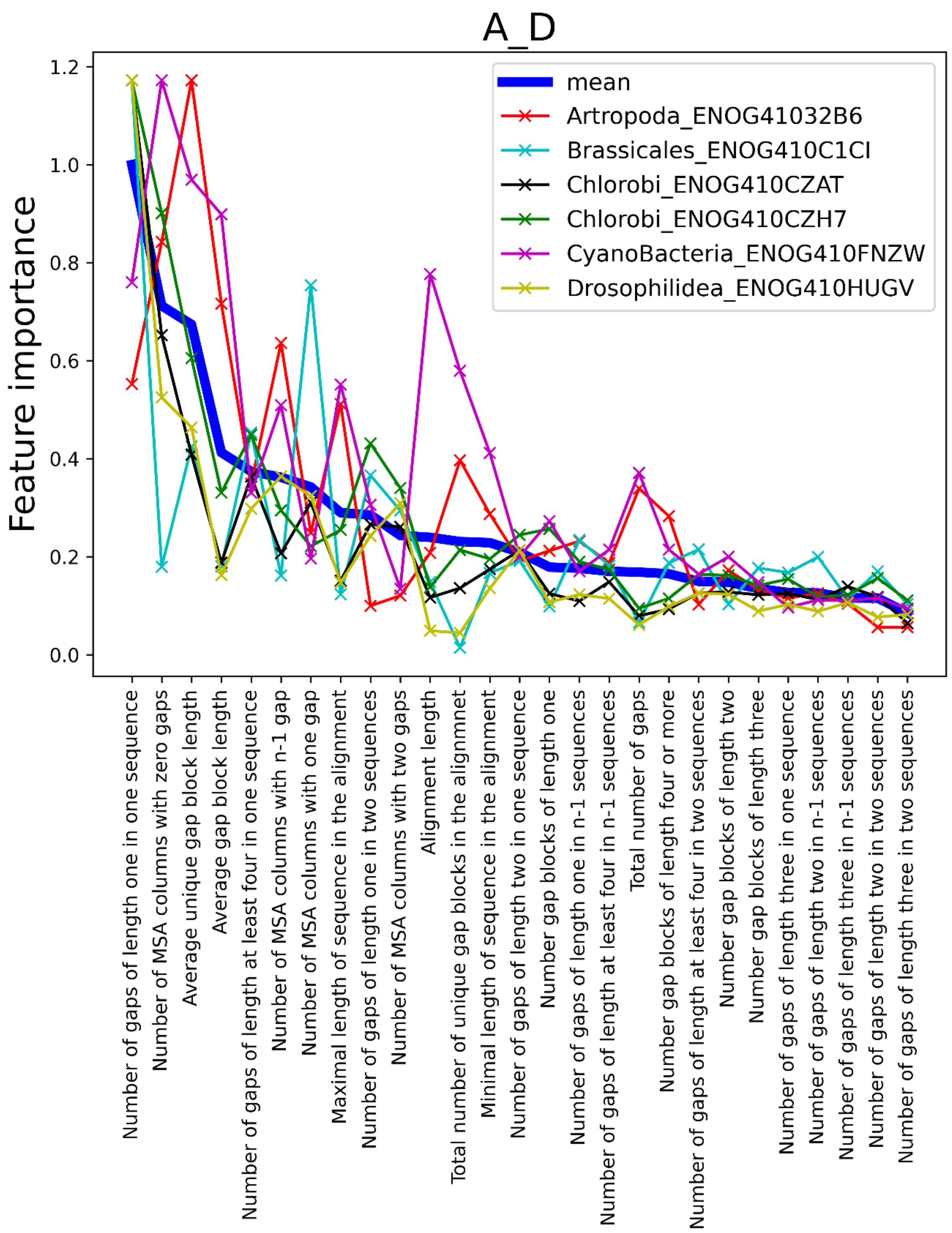

A_D
Feature importance
1.2
1.0
0.8
0.6
0.4
0.2
0.0
mean
Artropoda_ENOG41032B6
Brassicales_ENOG410C1CI
Chlorobi_ENOG410CZAT
Chlorobi_ENOG410CZH7
CyanoBacteria_ENOG410FNZW
Drosophilidea_ENOG410HUGV
Number of gaps of length one in one sequence
Number of MSA columns with zero gaps
Average unique gap block length
Average gap block length
Number of gaps of length at least four in one sequence
Number of MSA columns with n-1 gap
Number of MSA columns with one gap
Maximal length of sequence in the alignment
Number of gaps of length one in two sequences
Number of MSA columns with two gaps
Alignment length
Total number of unique gap blocks in the alignmnet
Minimal length of sequence in the alignment
Number of gaps of length two in one sequence
Number gap blocks of length one
Number of gaps of length one in n-1 sequences
Number of gaps of length at least four in n-1 sequences
Total number of gaps
Number gap blocks of length four or more
Number of gaps of length at least four in two sequences
Number gap blocks of length two
Number gap blocks of length three
Number of gaps of length three in one sequence
Number of gaps of length two in n-1 sequences
Number of gaps of length three in n-1 sequences
Number of gaps of length two in two sequences
Number of gaps of length three in two sequences

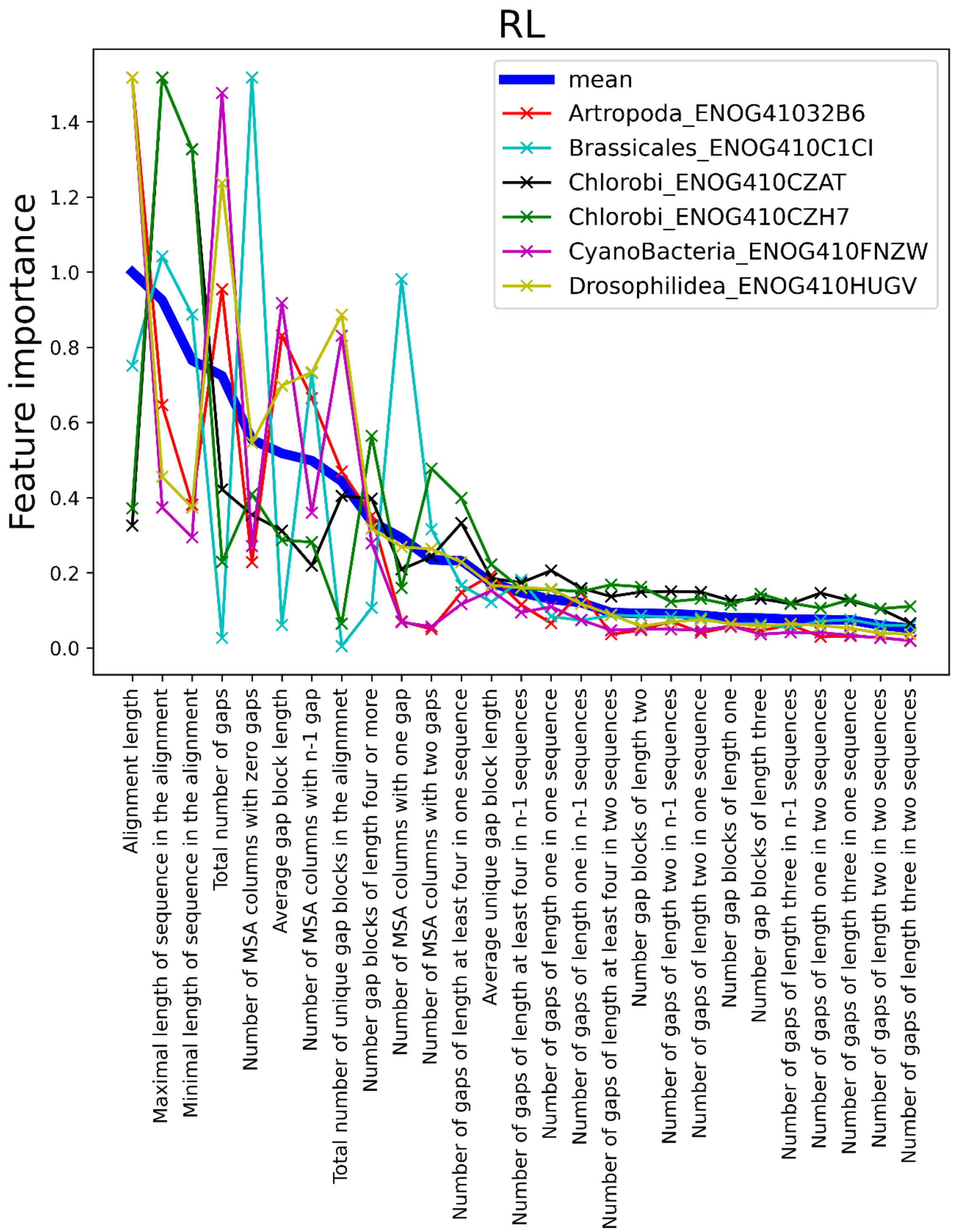


**Figure S2.** Each of the five panels corresponds to a different model parameter. Shown are the absolute values of the Lasso weight coefficients of each summary statistic, normalized by the maximum coefficient of the mean. Each thin line corresponds to one MSA from a specific phylogenetic group. The blue thick line in each panel represents the average over the six analyzed phylogenetic groups.

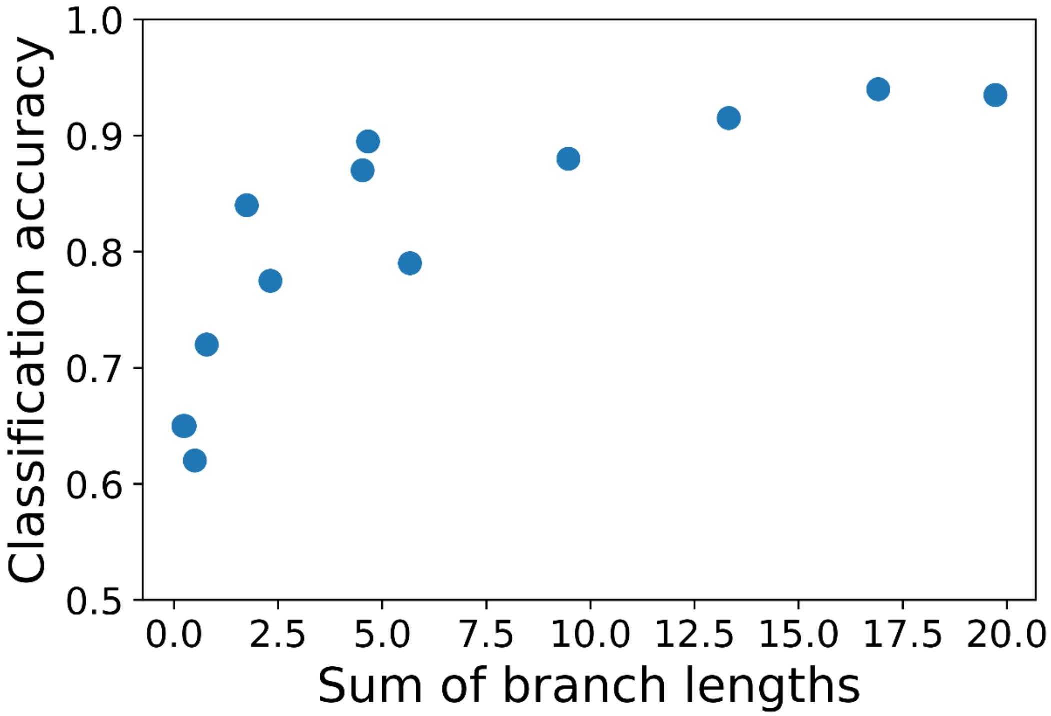


**Figure S3.** Neural network classification accuracy as a function of the total branch lengths.

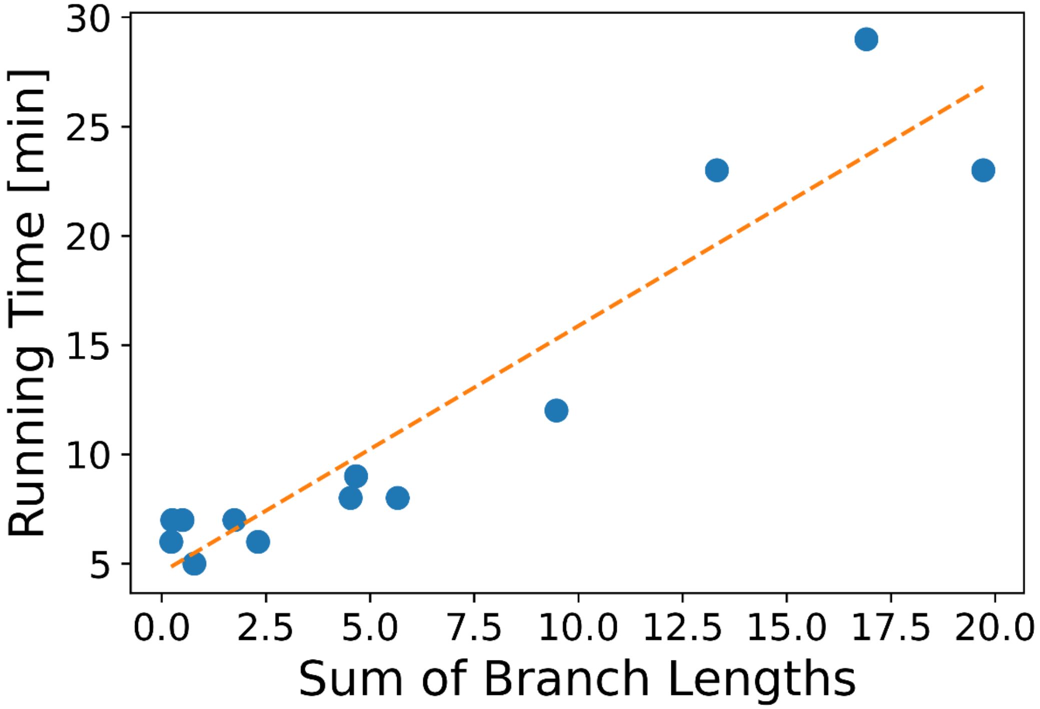


**Figure S4.** Running times are correlated to sum of branch lengths. $R^2 = 0.88$, P < 2e-6, $running\ time = 4.59\ \times sum\ of\ branch\ lengths + 1.13$. The sum of branch length for each accession number is given in Table S2. The data for this graph (EggNOG accession id, running times in minutes): (ENOG41032B6, 29), (ENOG410C1CI, 5), (ENOG410CZAT, 6), (ENOG410CZH7, 8), (ENOG410FNZW, 12), (ENOG410HUGV, 9), (ENOG410I2VE, 23), (ENOG410N476, 8), (ENOG410PK11, 23), (ENOG411668K, 7), (ENOG4116ES6, 7), (ENOG4118Z5J, 7), (ENOG41190IG, 6).

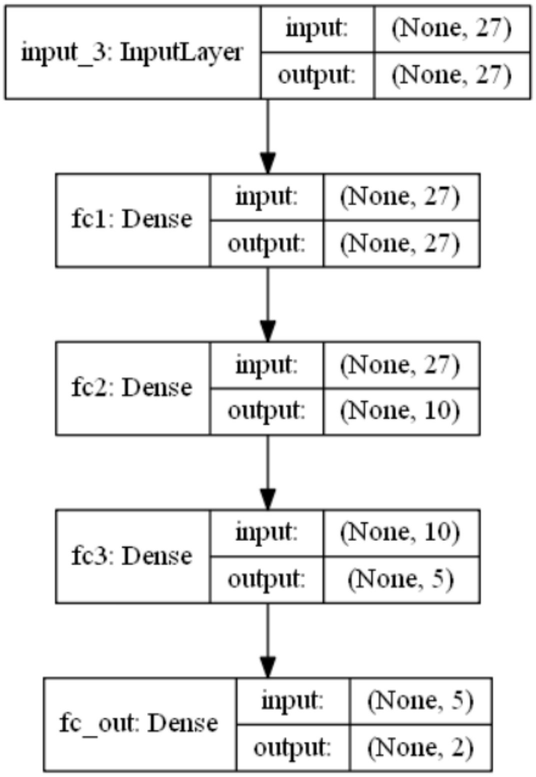


**Figure S5.** The architecture of the neural network for model selection. The 27 summary statistics normalized by their weights are the network input. The two hidden layers are fully connected ReLU layers. The last layer is a fully connected sigmoid layer with two outputs. The classification is determined by the maximal output.
